# Historical and cross-disciplinary trends in the biological and social sciences reveal an accelerating adoption of advanced analytics

**DOI:** 10.1101/2020.12.02.408989

**Authors:** Taylor Bolt, Jason S. Nomi, Danilo Bzdok, Lucina Q. Uddin

**Affiliations:** People Analytics, Deloitte Touche Tohmatsu Limited; Department of Psychology, University of Miami, Coral Gables, FL, USA; Department of Biomedical Engineering, McConnell Brain Imaging Centre (BIC), Montreal Neurological Institute (MNI), Faculty of Medicine, McGill University, Montreal, Canada; Mila - Quebec Artificial Intelligence Institute, Montreal, Canada

**Keywords:** data science, collaboration, statistics, machine learning, natural language processing, epistemology, multivariate statistics

## Abstract

Methods for data analysis in the biomedical, life and social sciences are developing at a rapid pace. At the same time, there is increasing concern that education in quantitative methods is failing to adequately prepare students for contemporary research. These trends have led to calls for educational reform to undergraduate and graduate quantitative research method curricula. We argue that such reform should be based on data-driven insights into within- and cross-disciplinary use of research methods. Our survey of peer-reviewed literature screened ∼3.5 million openly available research articles to monitor the cross-disciplinary usage of research methods in the past decade. We applied data-driven text-mining analyses to the methods and materials section of a large subset of this corpus to identify method trends shared across disciplines, as well as those unique to each discipline. As a whole, usage of *T*-test, analysis of variance, and other classical regression-based methods has declined in the published literature over the past 10 years. Machine-learning approaches, such as artificial neural networks, have seen a significant increase in the total share of scientific publications. We find unique groupings of research methods associated with each biomedical, life and social science discipline, such as the use of structural equation modeling in psychology, survival models in oncology, and manifold learning in ecology. We discuss the implications of these findings for education in statistics and research methods, as well as within- and cross-disciplinary collaboration.

## Introduction

Accelerated by trends in open-source science in the past decade, the methodological landscape of the biomedical, life and social (BLS) sciences is becoming increasingly complex. The classic statistical tools (e.g. *T*-tests, analysis of variance, other types of linear regression) taught in introductory statistics courses are nowadays perceived insufficient to prepare researchers for the age of big data, machine learning, and open source software. Concerned that BLS sciences educational training is failing to keep up with these trends, many researchers and statisticians have advocated for educational reform to introductory research methods and statistics courses (1–4). We argue that a crucial step in this direction is a more complete understanding of actual trends in method usage across BLS sciences. Such an understanding will offer valuable insights into the necessary methodological skills and knowledge needed to train early career scientists for future success in their disciplines and collaborations.

In this study, we conduct a systematic charting of research method usage across BLS disciplines over time. We applied natural language processing tools to a large corpus of open-access peer-reviewed literature (5). Our study aimed to map out the methodological landscape of the BLS disciplines and changing trends over the past decade. Here we use the term *research methods* to broadly denote any quantitative or qualitative method, algorithm, or metric used to describe, summarize or interpret a sample of data. ‘Study’ is also broadly defined as a peer-reviewed computer- or qualitative-based assessment of measured data points, including experimental, observational, or meta-analytic research. We carefully retraced trends in research methods across 1) time, and 2) research disciplines. From a temporal perspective, we identified research methods that have increased or decreased in prominence across BLS disciplines over the past decade (2009 to 2019). From a cross-disciplinary perspective, we identified research methods that are uniquely prominent within each BLS discipline, and the similarity or dissimilarity of BLS disciplines, in terms of their usage of research methods.

## Results

### Pre-processing and Analysis Summary

The primary goal of this study is to describe and understand usage shifts in research methods across BLS disciplines over time. We screened ∼3,500,000 articles published in a decade of research to accomplish this goal. We extracted mentions/adoptions of research methods from ‘Methods and Materials” sections of a large corpus of peer-reviewed articles (PubMed Open Access Subset; 5). We used a named entity recognition algorithm trained specifically for this purpose. We refer to these extracted mentions from the text as *method entities* – unique strings of alphanumeric characters that refer to a distinct research method. The extracted *method entities* then underwent a sequence of pre-processing steps including removal of unwanted characters and lemmatization (removing inflectional endings). The pre-processing workflow included a manual entity disambiguation step that classified *method entities* into more meaningful superordinate method categories. In addition to pre-preprocessing of method entities, articles were classified into a set of 15 research disciplines (see **Figure 1** and **Figure 2**) using a supervised machine learning framework pooling information from article titles, abstracts and journal names. The 15 disciplines were chosen by the authors from a survey of the corpus to balance breadth and specificity of the BLS literature. The entire pre-processing pipeline is illustrated in **Figure 1**.

**Figure 1.**
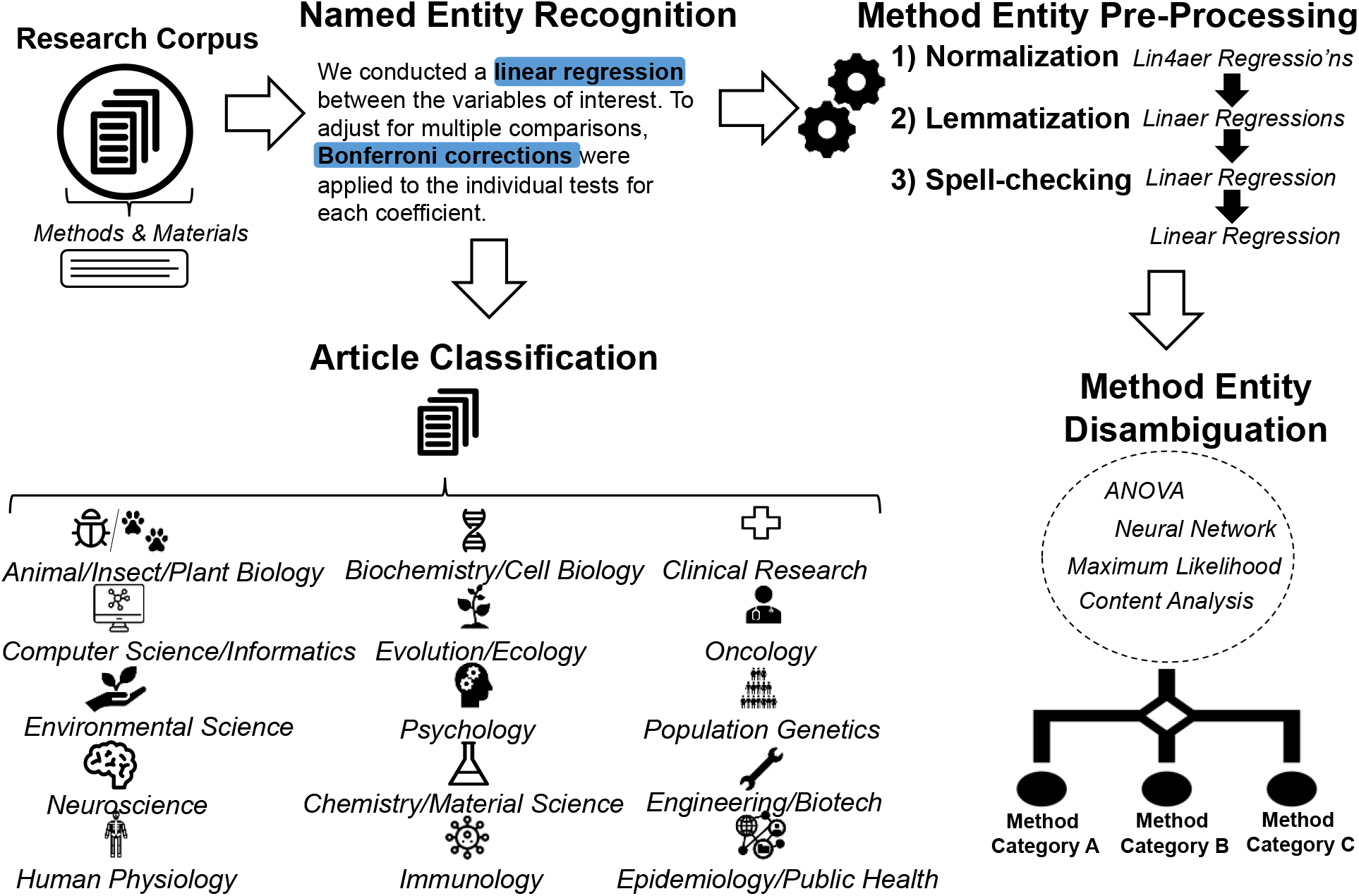
Pre-processing Pipeline. The pre-processing pipeline consists of 1) retrieval and parsing of full-text “Methods and Materials” sections, 2) named entity recognition of research *method entities*, 3) entity string pre-processing, 4) article classification into BLS disciplines, 5) and a manual entity disambiguation step whereby method entities are classified into superordinate categories of employed research methods (e.g. survival models, graph theory).

**Figure 2.**
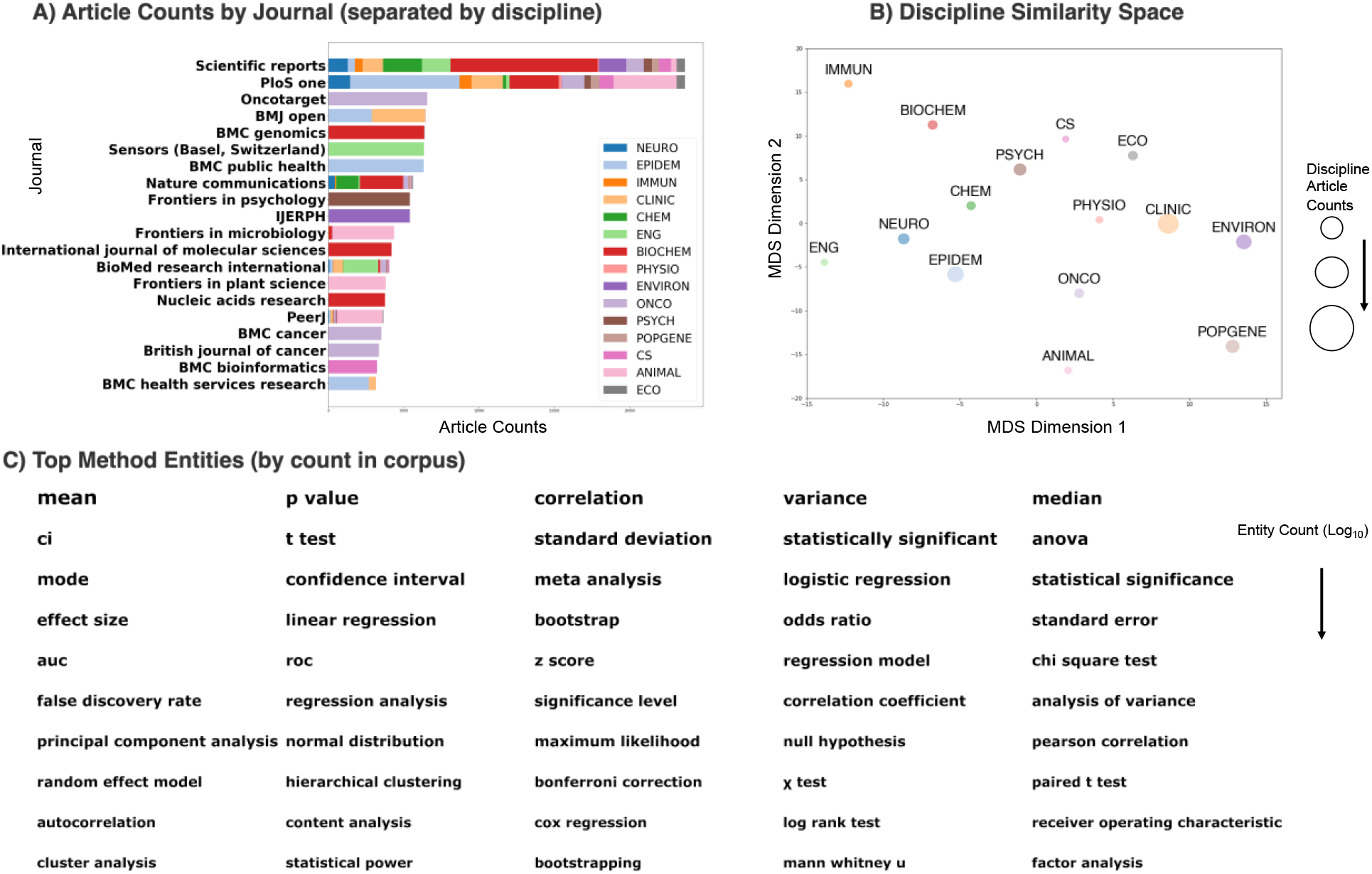
Journal, Discipline and Entity Statistics. (BIOCHEM: Biochemistry/Cellular Biology/Genetics; CLINIC: Clinical/Hospital Research; CS: Computer Science/Informatics; CHEM: Chemistry/Material Science; ENVIRON: Environmental/Earth Science; ECO: Evolution/Ecology; IMMUN: Immunology; ONCO: Oncology; PSYCH: Psychology; NEURO: Neuroscience; EPIDEM: Public Health/Epidemiology; ENG: Engineering/Biotechnology; PHYSIO: Human Physiology/Surgery; POPGENE: Population Genetics; ANIMAL: Animal/Insect/Plant Sciences). Translations of discipline abbreviations (e.g. CLINIC) are displayed at the top of the figure. **A)** A horizontal stacked bar plot displaying the number of articles for the top 20 journals in the corpus (defined in terms of article count). The percentage of articles per domain within a journal are proportionally shaded within each bar (IJERPH = *International Journal of Environmental Research and Public Health*). The research disciplines with the highest article counts were primarily biomedical and clinical disciplines. **B)** Multidimensional scaling plot displaying the similarity between research disciplines, in terms of total entity counts (summed across all articles in the discipline), on a two-dimensional space. **C)** Top 50 research method entity strings were sorted row-wise by the number of mentions across the corpus. The size of each entity string is proportional to the logged article count. The most frequently mentioned method entities were summary statistics and classical statistical methods (e.g., ANOVA, *T*-test, linear regression).

The method category counts per article (N_articles_ = 585,362) were used as input to two analytic pipelines: 1) *method usage trends* to observe temporal trends in research method usage over the time window of 2009 to 2019 (at an *annual* frequency), and 2) *discipline by research method probability analysis* to understand what research methods are unique to each BLS discipline. In addition, the pre-processed method entities before the entity disambiguation step were supplied to a third analytic pipeline: 3) *Analysis of Research Method Groupings* to discover data-driven clusters of research methods that frequently co-occur within and across BLS disciplines. To promote reproducibility and re-use, the full code for all pre-processing and analytic processes are provided on the following webpage: https://github.com/tsb46/stats_history.

### Journal, Discipline and Method Entity Statistics

The corpus of open-access peer-reviewed literature predominantly consisted of science-general journals, such as *Plos One, Scientific Reports, Nature Communications* (**Figure 1A)**. This observation highlights one advantage of the machine-learning classification of journal articles into scientific disciplines. The common practice of article classification by its journal publication would fail to capture the mixture of scientific disciplines contained within these science-general journals. Discipline-specific journals with high article counts included *Oncotarget* (Oncology), *BMJ Open* (Clinical Research, Public Health/Epidemiology), *BMC Genomics* (Biochemistry/Cellular Biology/Genetics), *Sensors* (Engineer/Biotechnology), *BMC Public Health* (Public Health/Epidemiology), and *Frontiers in Psychology* (Psychology). These discipline-specific journals publish peer-reviewed articles in a specific area of study and have a more focused readership. The disciplines with the highest article counts are primarily biomedical and clinical disciplines: Clinical Research (N=126,446), Biochemistry/Cellular Biology/Genetics (N = 76,607), Public Health/Epidemiology (N = 67,928), and Oncology (N = 52,487) (**Figure 1B**).

We deployed classical multidimensional scaling (MDS) to express each discipline’s total method entity counts in a parsimonious two-dimensional space (**Figure 1B**). The distances between the disciplines in the resulting plot reflect the dissimilarity/similarity in total research method counts. This approach made apparent that four disciplines stood out as *relative* outliers in research method usage: Oncology, Population Genetics, Evolution/Ecology and Chemistry/Material Sciences. As we observe below, among all 15 candidates, these select disciplines revealed a unique profile of research method usage. For illustration, we consider the discipline of Ecology/Evolutionary Sciences. Compared with other BLS disciplines, distance matrix and manifold learning methods (e.g., multidimensional scaling) are more widely used in the analysis of ecological data (6–8). Such methods have been found to be uniquely suited for the analysis of species composition and abundance data (9). For example, distance matrices constructed through metric/non-metric dissimilarity metrics (e.g., Bray-Curtis dissimilarity) are used to represent a species-by-sample/site matrix. Manifold learning methods are routinely used to analyze the resulting distance matrices (9). Manifold learning methods are often referred to as ‘ordination’ in ecology.

The most frequently mentioned method entities included summary statistics of a collection of data points (e.g., mean, variance, median, standard deviation), tools that are intimately related to statistical significance (e.g., *p* value, statistically significant, statistical significance, confidence interval), mean comparison tests (e.g., *T*-test, ANOVA), as well as (e.g., Pearson’s/Spearman’s) correlation coefficient and linear/logistic regression. Other frequently evoked method entities included dimension reduction and clustering techniques (e.g., principal component analysis and hierarchical clustering), resampling methods (e.g., bootstrapping), classification performance metrics (e.g., area under the curve, receiver operating characteristic), and survival models (e.g., cox regression). Interestingly, content analysis - a sometimes quantitative, sometimes qualitative coding method of documents to examine communication patterns – also appears in the top 50 most frequently mentioned method entities.

### Research Method Usage Trends

The methodological landscape of BLS sciences has seen a dramatic shift over the past decade. In order to track what research methods have increased or decreased in prominence over the past decade, we calculated the proportion of articles mentioning each method category (N = 126) per year (2009 – 2019) (**Figure 3**). As can be observed from **Figure 3** (top panel), classical statistical methods have remained the dominant analytic methods in the BLS sciences over the past decade. These classical methods include the ANOVA, *T-*test, linear regression, and chi square test. However, plotting these trends together on the same plot fails to visualize the relative increases/decreases in usage of these methods. When plotted *relative* to their own scale (i.e., different y-axis per method category), interesting trends appear. While still dominant, the widespread adoption of classical statistical methods has steadily weakened over the course of 2009 to 2019. Interestingly, the only classical statistical methods that we observed to gain in adoption over the past decade are effect sizes and confidence intervals. The increasing adoption of effect sizes and confidence intervals may reflect the increased pressure from institutions and researchers (10–12) to report effect sizes and confidence intervals along with significance tests in peer-reviewed research.

**Figure 3.**
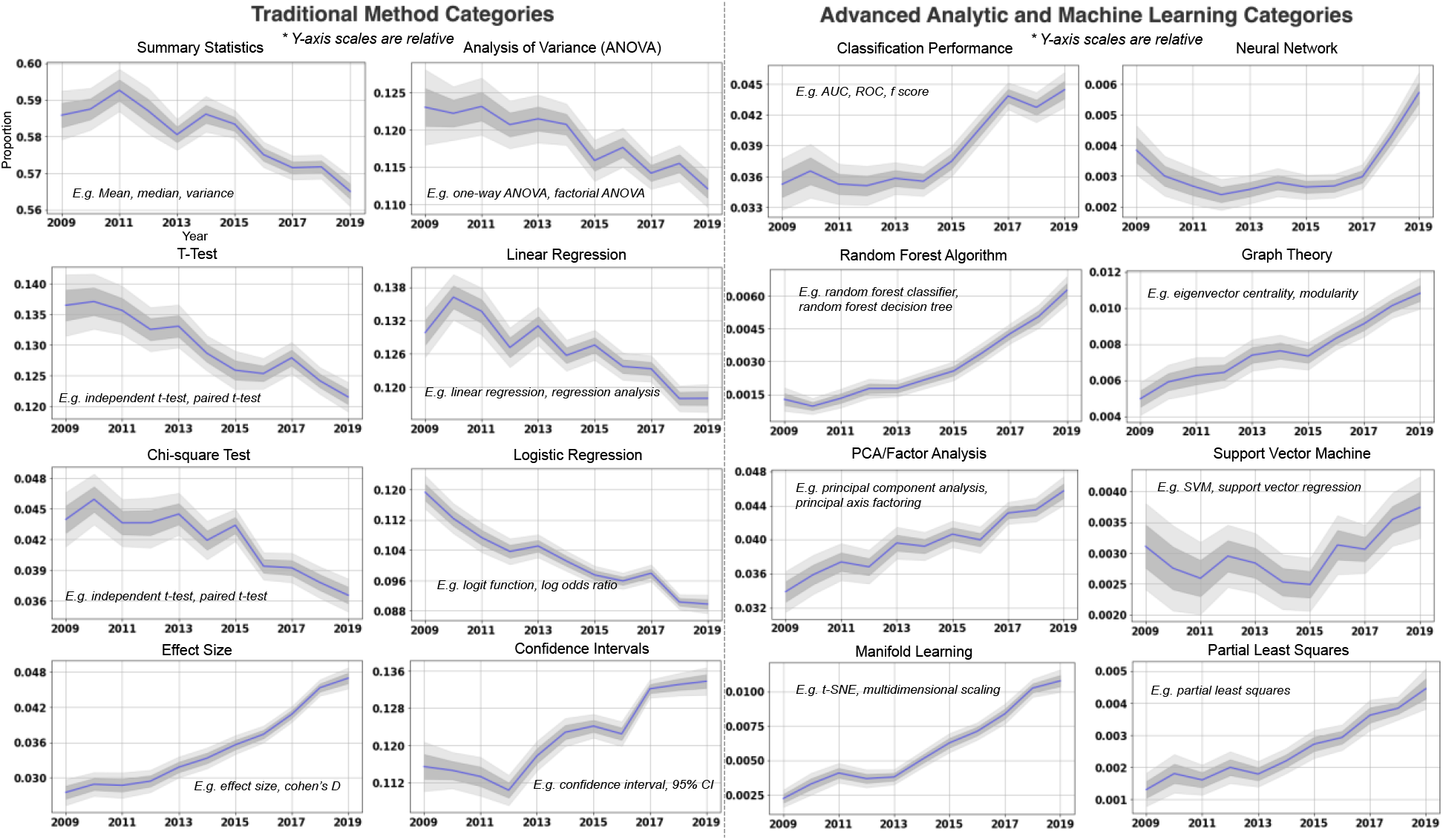
A Decade of Research Method Trends (2009-2019). Time-series of research method categories from 2009-2019 (annual frequency). For each research method category, the time-series represent the proportion of articles that contained an instance of that category per year. **Top Panel)** All time-series displayed in single plot with categories differentiated by color. Note that trends are difficult to discern due to base category proportions in the literature - ‘summary statistic’ entities appear in slightly over half (∼50%) of all articles. **Bottom Panel)** Time-series of individual categories (y-axis limits and scale relative to each category), separated into ‘traditional statistical method’ and ‘advanced analytic and machine learning’ categories. Random sampling variability for each proportion estimate was visualized using bootstrapped standard errors (SE) from 100 bootstrapped samples of articles at each time point (dark shaded region: ± 1 SE, light shaded region: ± 2 SE). Overall, advanced analytic methods have grown in adoption over the past decade. However, classical statistical methods have decreased in usage over the past decade.

Advanced statistical and machine learning methods have grown in adoption over the past decade. The method categories include metrics for evaluating classification performance (e.g., area under the curve, receiver operating characteristics), principal components analysis/factor analysis, random forest algorithms, support vector machines, manifold learning and partial least squares. In addition, the adoption of graph theoretical or network science approaches has grown in the BLS disciplines over the past decade. The frequency of usage of artificial neural networks in BLS disciplines remained relatively steady for the first half of the decade. However, a steep surge in adoption is observed after 2016, consistent with the recent explosion of interest in ‘deep learning’ (13, 14).

Other recent developments in research method usage are also worthy of note (**Figure 4**). First, the adoption of qualitative research methods has steadily grown in the BLS sciences over the past decade. These methods include thematic analysis (15), framework analysis (16), content analysis (17; though this method is sometimes used in quantitative manner - e.g., word counts), and grounded theory methodology (18). The common features of these qualitative methods are their application to unstructured or non-relational data (e.g. text, audio or video recordings) - an exponentially growing source of data in both industry and the BLS sciences (19–21).

**Figure 4.**
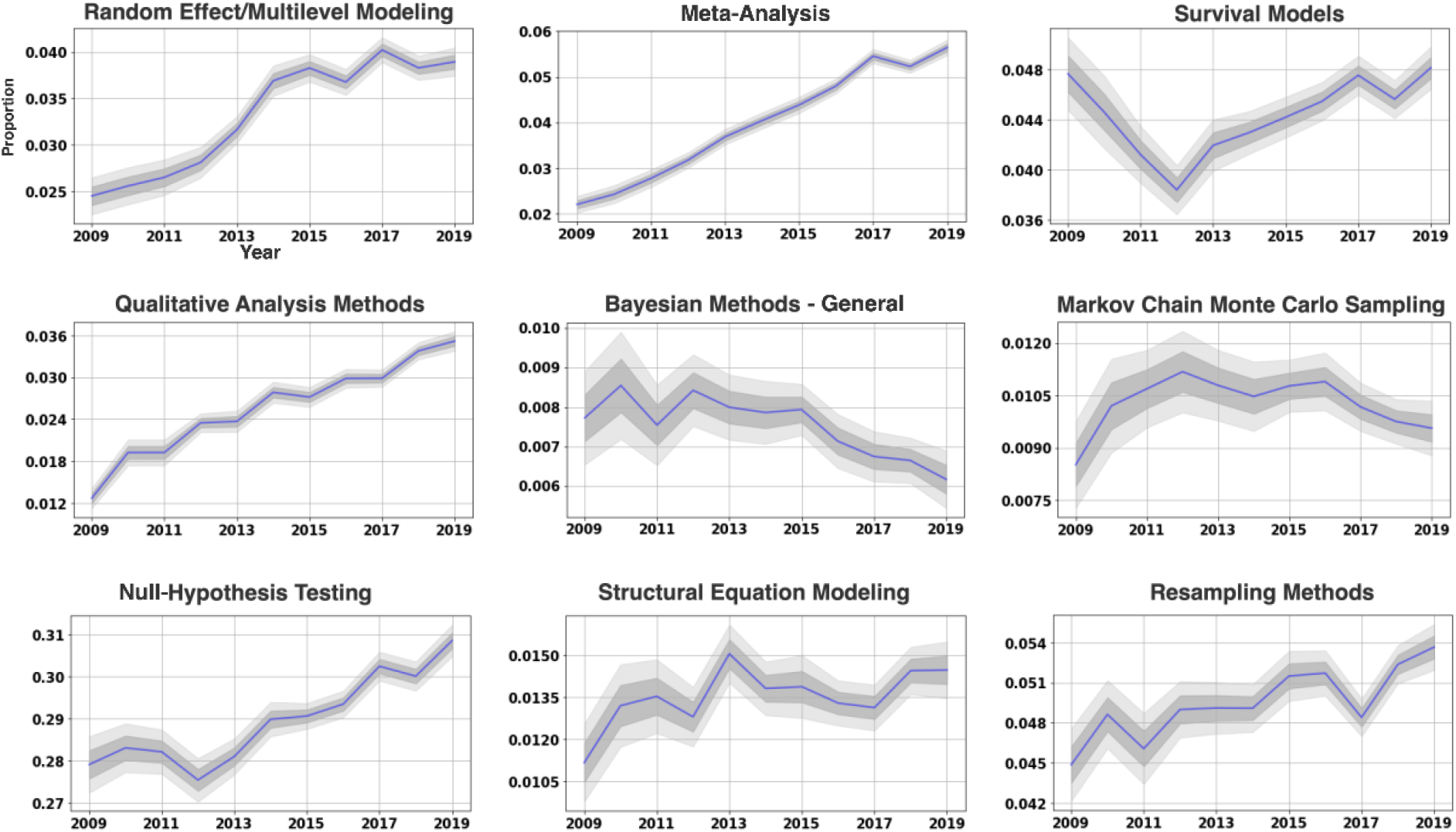
Additional Research Method Trends. Additional time-series of research method categories from 2009-2019 (annual frequency). For each research method category, the time-series represent the proportion of articles that contained an instance of that category per year. Random sampling variability for each proportion estimate was visualized using bootstrapped standard errors (SE) from 100 bootstrapped samples of articles at each time point (dark shaded region: ± 1 SE, light shaded region: ± 2 SE). Qualitative methods, meta-analysis, and null-hypothesis significance testing have steadily grown in adoption over the past decade. Other research method trends exhibit more complex patterns – e.g. survival models exhibited a steady decrease in the first half of the decade, followed by a steady increase in the latter half.

Another noteworthy trend of interest is the lack of growth in adoption of Bayesian methods for the BLS sciences. Bayesian methods include general Bayesian concepts (e.g. prior distribution, posterior distribution, Bayesian estimation), as well as sampling methods such as Markov chain Monte Carlo (MCMC) sampling. In contrast, null-hypothesis testing methods and concepts (e.g., *p* value, statistical significance, null hypothesis, alternative hypothesis) have exhibited a steady increase in application in the BLS sciences over the past decade. This lack of enthusiasm for Bayesian methods may be somewhat surprising, given the increasing advocacy of these methods as an alternative to the predominant use of null-hypothesis testing, with its perceived flaws (22, 22–24).

### Unique Method Usage by Discipline

To examine method categories uniquely associated with each BLS discipline, we used a contingency table approach. Specifically, we modeled the probability that an article belongs to each discipline vs. the rest (i.e., binary dependent variable) as a function of the research method category mentions in that article. **Figure 5** displays the unique set of method categories most frequently used in that discipline. For example, Gaussian Process regression (GPR; known as ‘kriging’ in geostatistics), employed as a method for spatial smoothing by interpolation, is uniquely associated with Earth/Environmental sciences. GPR is uniquely suited for the often required need to infer the level of quantities (e.g., minerals) at spatial locations for which no or sparse data were measured. Another example is the uniquely predominant use of partial least squares in chemistry, or chemometrics. PLS, a multivariate technique that predicts a set of response variables (Y) based on a set of predictor variables (X), is often used in chemometrics to relate properties of chemical samples (e.g. spectral properties) to their chemical composition (e.g., sample concentrations) (25). Other discipline-method groupings include structural equation modeling in psychology, independent component analysis in neuroscience, meta-analysis in clinical/health research, and survival models in oncology.

**Figure 5.**
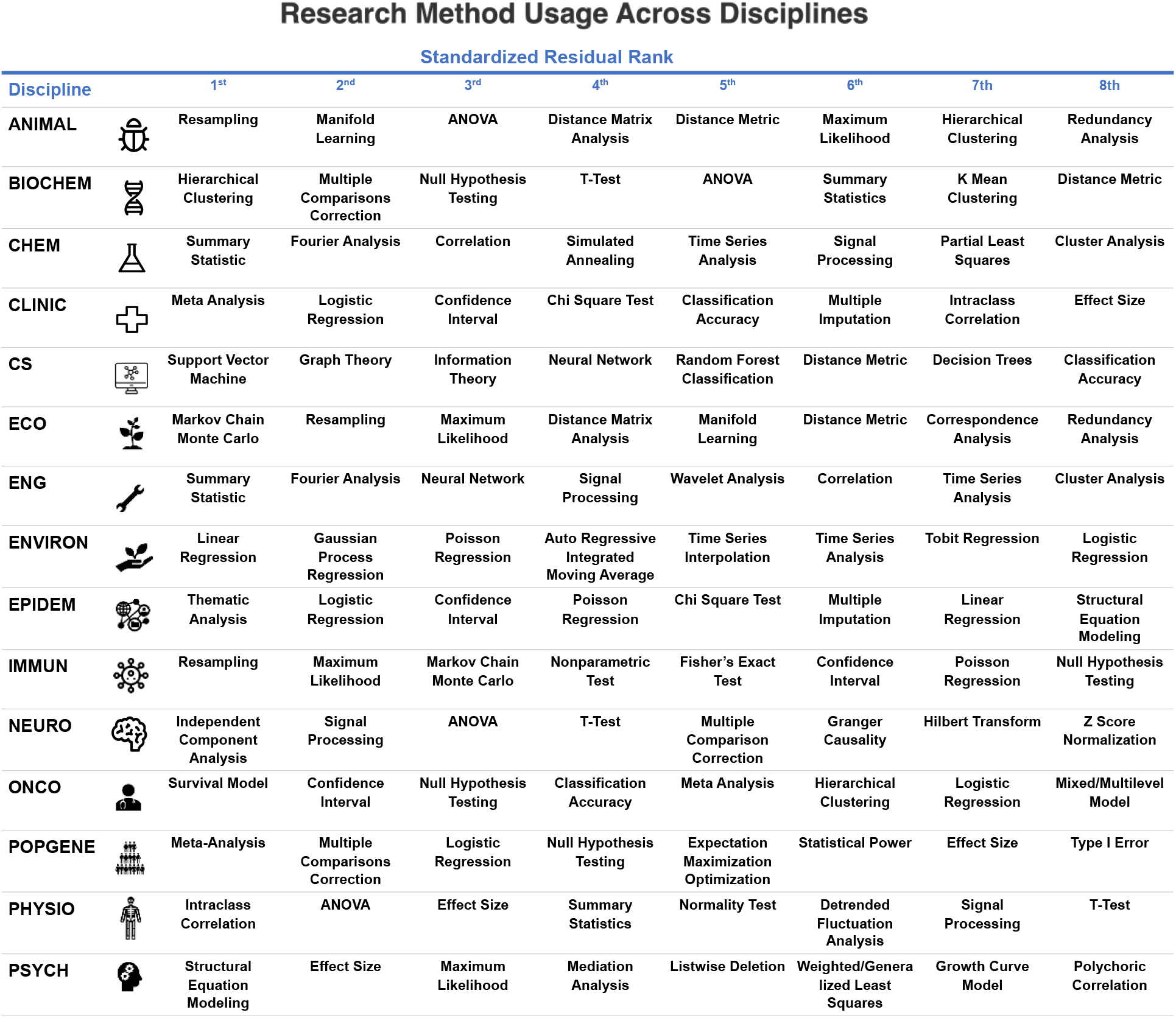
Research Method Usage Across Disciplines. Top eight standardized chi-square residuals for each discipline from the contingency table analysis. The greater the chi-square residual, the greater the difference between the observed and expected number of method entities within that discipline. Method entities are arranged by ranking from left to right. Disciplines have a unique set of research methods frequently applied to their subject matter: e.g. information theory in computer science, independent component analysis in neuroscience, and meta-analysis in population/behavioral genetics.

### Research Method Groupings

Research methods in the BLS sciences are rarely used in isolation. Rather, a set of methods are applied jointly, or in sequence, to understand a dataset. We term these frequently co-occurring research methods, ‘method groupings’. To directly extract coherent constellations of methods and understand how they vary across BLS disciplines, we applied a tensor decomposition approach to a method entity (N = 1,218) co-occurrence by discipline (N = 15) tensor (**Figure 6**; top left panel). For illustration, we displayed the selected components for a 20 component (i.e., 20 rank-one tensors) solution (**Figure 6**). Each component is associated with separate weights for method entities and disciplines, indicating the method entities and disciplines most associated with the component, respectively.

**Figure 6.**
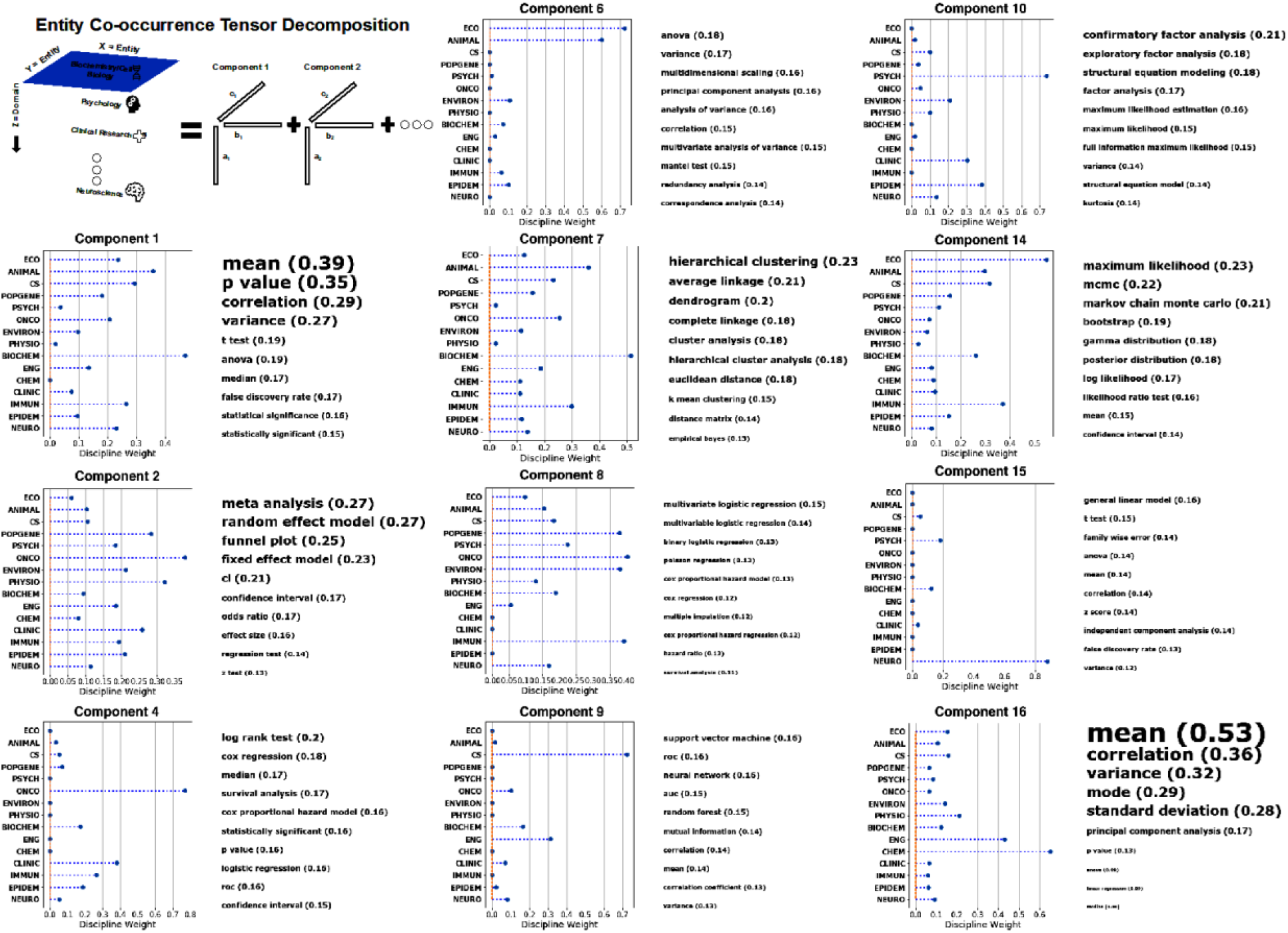
Co-occurrence Patterns in Method Usage Across BLS Disciplines. To understand what research methods are frequently used together in the same study, we conducted a tensor decomposition of a method co-occurrence by discipline tensor. The tensor decomposition analysis simultaneously models the co-occurrence between research methods, as well as their frequency of usage in each discipline. This figure displays the discipline and method entity weights from the tensor decomposition analysis. Components from the tensor decomposition are referred to as ‘method groupings’, or groups of research methods that frequently occur together in study method sections. The top left panel provides a visual illustration of the tensor decomposition (non-negative CANDECOMP decomposition) of the entity co-occurrence by discipline tensor. The first two dimensions of the tensor represent the logged sum of co-occurrences between each pair of research methods. The third dimension splits out the research method co-occurrences by discipline (i.e., the research method co-occurrences of articles within each discipline). For each component, or ‘method grouping’, a stem plot illustrates the weights for each discipline, as well as the top 5 method entity strings, in terms of their weights (sized by their weight). For each component, the discipline weights represent the frequency of usage of that component across each discipline. Some sets of methods are represented across all BLS disciplines (e.g., component 2), while others are concentrated within one or two disciplines (e.g., component 15).

The method groupings revealed by the tensor decomposition can be roughly classified into cross-discipline and within-discipline categories. Cross-discipline method families include components with a broad representation across the BLS disciplines (i.e., relatively more even distribution of discipline weights). For example, components 1 and 16 had non-zero weights for the majority of BLS disciplines. These components had high weights for both summary statistics and null-hypothesis testing methods. Interestingly, component 1 had higher weights for null-hypothesis testing concepts (e.g., *p* value, statistical significance) compared to component 16. In addition, component 1 was more associated with life science disciplines. Instead, component 16 was more associated with the engineering, chemistry and material science disciplines. Another cross-discipline method grouping was component 7: a group of unsupervised cluster analysis methods/metrics - primarily hierarchical clustering (e.g., average linkage, dendrogram). This component was most associated with the biochemical, animal/plant and immunological disciplines. Another cross-discipline method grouping was component 8: a group of regression approaches that represent particular instances of generalized linear models, including output distributions following logistic, Poisson, and truncated (e.g., Cox regression) laws. This component was most associated with the categories of population genetics, oncology, earth/environmental sciences and immunology disciplines.

We subsequently focused attention on the discipline-specific method groupings: method groupings with almost exclusive use in one or a small subset of BLS disciplines. One example was component 6, a method grouping with exclusive use in ecology and animal/plant/insect disciplines. This set of methods included ANOVA-based methods, manifold learning (e.g., multidimensional scaling), and distance matrix analyses (e.g., Mantel test). As noted above, manifold learning and distance-matrix methods are uniquely suited to analyses of species composition and other types of data regularly collected in these disciplines. The appearance of ANOVA-based methods in this seemingly unrelated group of methods may seem surprising, but owes to the fact that variance partitioning of *distance matrices* is a historically common practice in the ecological disciplines (26). Another, perhaps surprising, discipline-specific method grouping is component 15: a set of methods/concepts, including the general linear model, *T*-tests and family-wise error correction. This method grouping belonged exclusively to the discipline of neuroscience. This neuroscience-specific method grouping owes its existence to the development of statistical parametric mapping of neuroimaging data developed in the late 1990’s and early 2000’s, which rely heavily on the concept of a general linear model and random field theory for family-wise error correction of statistical maps (27, 28).

## Discussion

It was famously stated that “Data science will become the sexiest job in the 21^st^ century” (Hal Varian, Google). We offer an automated 10-year survey of ∼3.5 million open research papers. This study aimed to detect and expose the trajectory of usage patterns that characterize distinct BLS disciplines. Our grass-roots approach provides a sociological snapshot of the ongoing methodological shifts in a variety of scientific communities. We find that multivariate algorithms have become rapidly embraced in response to the expanding data deluge. Our results provide valuable pointers for how university curricula should be revised to meet the urgent need for training a new generation of quantitatively literate scientists.

‘Multivariate and machine learning approaches’ is a broad label, referring to a wide variety of quantitative methods. These methods are often taught in advanced statistics and computer science courses: principal component analysis, partial least squares, support vector machines, random forest classification algorithms, graph theory and artificial neural networks. While some of these methods are quite old (e.g., principal component analysis was first developed in the early 20^th^ century), others are relatively new and still developing (e.g., artificial neural networks have only seen broad use in the past decade). The increase in adoption of advanced analytic methods could be due to several reasons: 1) the collection of larger and more complex datasets, 2) the recent popularity of data science as a tool in academia and industry, or 3) an increasing realization among researchers that manuscripts containing advanced analytics are more likely to impress reviewers and editors. Either way, our findings reinforce the concern that statistics and research method education in the BLS sciences is falling behind and struggling to ‘keep up’ with the rapid pace of contemporary research in the age of big data, machine learning and open source software (2, 3). To prepare future practitioners in their disciplines, introductory research methods and statistics courses in the BLS sciences may need to be reimagined around a ‘data science’ focus (4, 29).

Our survey demonstrates that research methods for data analysis can vary widely across BLS disciplines. Several explanations can be offered for the distinct usage of data analysis methods between BLS disciplines. Perhaps the primary driver of a discipline’s adoption of research methods is the simple observation that the subject matter lends itself to the assumptions and goals of select research methods. For example, consider the observed disproportionate use of structural equation modeling in the discipline of psychology (**Figure 5** and **Figure 6** – Component 10). Psychological research, since the advent of cognitive psychology (30), routinely relates observable behavior such as task performance or questionnaire responses to unobserved or latent variables. The desire to explore causal structure among these latent variables has led to the systematic adoption of structural equation modeling – a technique to specify and test causal structures among latent and observable variables (31, 32). Similar explanations can be offered for other method-discipline pairs such as survival models and oncology. Other differences may arise from historical contingency, with no necessary connection between an analysis method and the subject matter it is applied to. For example, consider the predominant use of Fisher’s Exact Test in immunology vs. the Chi-square Test in clinical research (**Figure 5**). Both are statistical significance tests of the association between two categorical variables. The appropriate context for each test is controversial among statisticians (33, 34). Despite this controversy, our analysis indicates Fisher’s Exact Test is generally preferred over the chi-square test in the field of immunology, and vice versa in clinical research. Thus, the usage of one research method over another in a BLS discipline can be due to principled statistical reasons, and sociological or historical contingency.

Differences in analytic method usage have concrete implications for the direction of research in each BLS discipline. The choice of experimental/observational design often entails the subsequent analytic method used to analyze the data, but a reverse influence occurs as well: the researcher’s knowledge of available analytic methods informs their experimental or observational design. For example, ANOVA models for analysis of group means have a historically close relationship with experimental design in social and life science research (35). In other words, the influence between the choice of data analysis method and how data is collected operates in both directions. This observation underlies the potential for cross-fertilization and mutual inspiration between BLS disciplines by the discovery of new methods for data analysis, as well as novel ideas around data collection. While many advocates of cross-disciplinary collaboration have emphasized the joining together of different theoretical and subject-matter expertise (36), our findings emphasize a further methodological benefit of collaboration, which affords practitioners access to novel methods of data analysis not widely known in their own disciplines.

It should be noted that the corpus used in this analysis is limited in many respects. First, it only contains open-access articles made available by an open-access journal or an NIH-funded author. Thus, a sizable collection of peer-reviewed research in the past decade is systematically missing from this analysis. However, we assume that the type of publisher - open-access or subscription-based – is not a significant determiner of the methods used within a discipline.

Second, some scientific disciplines are missing from this survey, including experimental and theoretical physics, anthropology, astronomy, cosmology, economics, sociology, and geology. Future studies with a more comprehensive corpus of scientific publications will provide deeper insight into the historical and cross-disciplinary trends in scientific data analysis.

Our survey of peer-reviewed literature reveals the rapid pace of change in research methods in as few as 10 years. A comparable pace of change will be required in education of budding scientists. Equally important is the observed analytic diversity of BLS sciences of the past 10 years. The diverse analytic toolsets across BLS disciplines promises large pay-offs for cross-disciplinary collaboration. In this vein, the recent advent of ‘big data’ and open-source science is at least as much an opportunity for adequately training the next generation of researchers, as a challenge.

## Methods

### Peer-Reviewed Literature Corpus

Two sources of peer-reviewed literature were used for this analysis: the Pubmed Central Open Access Subset (PMC OAS) (N=2,869,889 articles at time of study); and the Pubmed Central Author Manuscript (PMC AM) collection (N=659,133 articles). The PMC OAS provides access to full-texts from a total of 14,722 open access peer-reviewed journals (at time of study). The PMC AM collection provides access to full texts of manuscripts made available in PMC by authors in compliance with the NIH Public Access Policy. Both sources form part of PMC’s open access collection (37) (https://www.ncbi.nlm.nih.gov/pmc/tools/textmining/). Bulk downloads of the full OAS and AM collection articles were conducted using the PMC FTP service. Overall, a total of 3.5 million articles were downloaded and screened for our analysis.

### Article Parsing and Method Section Identification

Both OAS and AM text corpora are openly available in a structured and standardized form to the public on PMC’s webpage. Both corpora were downloaded in XML format. Each XML article file is organized into article metadata and article text separated into article sections (e.g., Introduction, Methods & Materials, Results, etc.). As the primary interest of this study was exploring usage patterns of research methods in this pool of articles, we retrieved sections of the XML articles that correspond to the methodology section of the article (e.g., ‘Methods and Materials’). Given that methodology sections have no standardized title, we pulled any sections of text from each XML article that contained in any of the following sequence of strings: ‘method’, ‘material’, ‘measure’, ‘analysis’, and ‘statistical’. As we were interested in *original studies*, XML articles not containing any of these section search strings were excluded from analysis, such as literature reviews, book reviews, commentaries, etc. Of the total XML articles in the peer-reviewed literature corpus, N_text_ = 1,276,452 articles with the above section search strings were retrieved.

### Named Entity Recognition of Research Method Entities

In the resulting corpus of 1,276,452 methodology section texts, it would be infeasible to manually tag each research method phrase in every section text. Thus, we opted for an automated recognition method for the purpose of detecting research method phrases. We used a combined phrase-matching/rule-based and machine-learning approach. First, we developed a large list of phrases (N_phrase_=700) corresponding to commonly used research methods across BLS disciplines. These were used as a rule-based matching approach to detect research method phrases in the section texts. To ensure that the discovered research method phrases were not biased towards our list of pre-specified phrases, we selected 2,625 random method section texts with tagged phrases from our rule-based approach and fed them as training examples to a statistical named entity recognition (NER) algorithm. The objective of the NER algorithm is to utilize the rule-based training samples as context ‘clues’ for detecting research method phrases more generally (i.e., those outside the original phrase list). To perform NER on our full corpus, we used the convolutional neural network (CNN) algorithm provided by the open-source spaCy python package (38), with standard parameters for training (100 *iterations*, 0.2 *dropout*, mini-batch training). The trained NER model was applied to the entire corpus of methodology section texts to generate the final list of detected research method phrases, or entities. Of the total corpus of 1,276,452 methodology section texts, at least one research method entity was detected in approximately half of the texts (N_text_=662,482). In the main text, we refer to the detected research method phrases from the trained NER model as *method entities*. The total number of unique method entities (before pre-preprocessing) discovered from the NER algorithm was N_entity_=16,020.

### Research Method Entity Pre-processing

The NER algorithm yielded a unique list of method entities, which was represented as a distinct sequence of characters. Different entities (i.e., sequence of characters) can refer to the same research method: for example, one could refer to a *T*-test as ‘T-tests’, ‘T test’, ‘Ttest’, etc. To attempt to correct for small spelling differences such as these, each method entity string underwent a sequence of pre-processing steps for harmonization: 1) lowercase characters, 2) removal of non-alphanumeric or non-Greek characters (e.g., hyphenations, quotes, commas), 3) lemmatization of words of the tokenized (i.e., separated into words) entity strings, and 4) converting entity strings that occur rarely (N < 5) to more commonly used spellings (within a max difference of two characters) using the SymSpell algorithm implemented in: https://github.com/wolfgarbe/symspell. As another measure of quality control, only pre-processed entities with a minimum of 10 occurrences were included in the final method entity vocabulary. The *final method entity vocabulary count* after pre-processing was N_entity_=1,218. Examination of journal article counts revealed that *PloS One* contained a disproportionate number of articles in the corpus (N ∼ 80,000). The the next largest journal – *Scientific Reports* – contained N ∼ 23,000 articles. To counteract this potential bias from the data, we only included a random subset of *Plos One* articles in the corpus (N ∼23,000 – number of the next highest journal count). After pre-processing, the final pool of articles was N_text_= 585,362.

Before the 1) *trends analysis* and 2) *discipline by method probability analysis*, we manually disambiguated method entities belonging to the same class of methods. Many method entities in the final vocabulary were either the same method with different references (e.g., ‘cox regression’ and ‘cox proportional hazards regression’), or belonged to a more directly meaningful overarching category of analytic methods (e.g., ‘one-way analysis of variance’ and ‘factorial analysis of variance’ – ANOVA methods, or ‘modularity’ and ‘betweenness centrality’ – graph theory). Thus, we manually classified each of the method entities denoting research methods (N = 1,218) into more parsimonious (superordinate) categories for understanding temporal and across-discipline trends in research method usage. We refer to these superordinate categories of research methods as *method categories*. Method entities were grouped together in a method category if their underlying mathematical/statistical models were similar (e.g., ‘independent t-test’ and ‘paired t-test’ as belonging to *T*-test), or belonged to a commonly grouped class of models/metrics (e.g., ‘survival models’ or ‘structural equation models’). The total number of method categories resulting from our manual classification was 126. However, of the 1,218 method entities in the final vocabulary, 413 method entities were unclassified due to ambiguity (e.g., ‘test score’). The 3) *method grouping analysis* (see below) used the pre-processed method entities (N = 1,218) before the entity disambiguation step. This was because the *method grouping analysis* aimed at a data-driven grouping of frequently co-occurring research methods, as opposed to a categorization based on mathematical similarity or convention.

### Article Classification into Research Disciplines

One central objective of this study is to understand *cross-disciplinary* usage in research methods. This objective requires that the articles in the corpus are first classified into separable disciplines (e.g., biochemistry, epidemiology, psychology, etc.). A simple approach would be to manually classify each journal (or publication) in our corpus into a discipline, such that each article belonging to that journal would be classified with that discipline. While a much more manageable task than manually classifying each article, this approach runs into two problems: 1) not all articles in a domain-specific journal (e.g., PLOS Biology, Journal of Neuroscience, EMBO Journal) can be easily classified into one BLS discipline and 2) domain-general journals (e.g., PLOS ONE, Nature Communications, Science) cannot be classified into a single BLS discipline. Thus, we chose to use a machine-learning text classification approach to classify each *single article* into a set of pre-specified BLS disciplines. This approach allowed for more flexible classification at the level of each journal article - allowing for different article classifications within the same journal.

We chose a set of 15 BLS discipline categories for article classification: *Biochemistry/Cellular Biology/Genetics* (BIOCHEM), *Clinical/Hospital Research* (CLINIC), *Computer Science/Informatics* (CS), *Chemistry/Material Sciences* (CHEM), *Environmental/Earth Science* (ENVIRON), *Evolution/Ecology* (ECO), *Immunology* (IMMUN), *Oncology* (ONCO), *Psychology* (PSYCH), *Neuroscience* (NEURO), *Public Health/Epidemiology* (EPIDEM), *Engineering/Biotechnology* (ENG), *Human Physiology/Surgery* (PHYSIO), and *Population Genetics* (POPGENE). Importantly, *we make no claim that this categorization represents the most optimal division of BLS disciplines* – a potentially infinite number of categorizations could be more/less useful in certain contexts and overlap in research topics between the categories of any division will be prevalent. Rather, we make the claim that this categorization provides a useful/pragmatic division of BLS disciplines given the distribution of publications in our corpus. To classify articles in the corpus into the 15 BLS disciplines, we input a bag-of-words feature-set (1-gram, 2-gram and 3-gram tokens), generated from the article abstract, title, and journal title, to a multinomial naïve Bayes (MNB) classification algorithm. We utilized a version of the MNB algorithm that corrects for unequal number of instances across categories (39), as was present in our corpus. To reduce the number of features input to the MNB algorithm and improve prediction accuracy we applied a chi-square feature selection approach. Specifically, we chose the top 15,000 features from the bag-of-word feature-set, where each feature was sorted by their chi-square statistic value with the 15 discipline categories. The Chi-square Statistic is a measure of dependence between two categorical variables. The final MNB model for training involved 15,000 features, 15 discipline categories to be predicted, and 1,470 training samples. We assessed the model accuracy using a repeated K-fold cross-validation approach (#folds = 10, # of repeats = 10). The final MNB model achieved an accuracy of 74.9%. We found that the MNB approach performed better than other classification approaches (e.g., random forest decision tree classification, support vector machines, and logistic regression) on our dataset, in terms of classification accuracy.

### Discipline Similarity Analysis

To examine the similarity of research method usage between disciplines, we used a multidimensional scaling approach. First, we summed the counts of all method entities (N = 1,218) per discipline. We then computed the Euclidean distance between all pairs of z-score normalized count vectors (one vector per discipline). The distances were then projected to a 2-dimensional space using a classical multidimensional scaling algorithm.

### Method Trend Analysis

In order to understand temporal changes in research method usage across BLS sciences, we computed annual counts of method categories starting at the beginning of 2009 to the end of 2019 (N = 11 time points). We chose an annual time-resolution for the following reasons: 1) some journals in our corpus published at a limited time-frequency (e.g., annually or bi-annually) and 2) it was reasoned that large-scale trends in research method usage within a discipline occur at a low-frequency (i.e., over years, as opposed to months). The annual article count in our corpus increased at approximately a steady linear frequency from 2009 to 2019: 2009 - *21,440*, 2010 – *25,978*, 2011 – *32,141*, 2012 – *40,362*, 2013 – *54,184*, 2014 – *60,173*, 2015 – *70,513*, 2016 – *78,966*, 2017 – *83,337*, 2018 – *87,361*, 2019 – *64,492*. To account for this increase in article frequency, we normalized the raw frequency counts of each method category by the total article counts per year – i.e., we computed the proportion of counts of each method category to the total article counts per year. Thus, all method category trends are displayed as proportions of the total number of articles per year. To estimate the statistical accuracy of the original proportions for a given method category and year, we obtained an estimate of the sampling variation by constructing 100 bootstrapped samples of all articles within a year and re-calculated the proportion by total article count for that method category. Standard errors were estimated from the standard deviation across the 100 bootstrapped proportions and displayed as shaded regions around the observed proportion trends.

### Discipline by Research Method Probability Analysis

In order to understand the unique clustering of method entities between disciplines, we used a chi-square contingency table approach. This approach provides a straightforward way to identify those categories of research methods that are used more frequently in a discipline relative to others. Specifically, we organized the data into a discipline by method category count matrix and calculated the Pearson chi-square standardized residual for each cell (i.e., discipline and method pair). The standardized residual is simply the Pearson residual (observed – expected frequency) divided by the square root of the expected frequency. The larger the positive value of the standardized residual, the greater than expected number of articles in a BLS discipline mentioning that method category (in its ‘Materials and Methods’ section). The top eight standardized residuals for each BLS discipline are displayed in **Figure 5**.

### Analysis of Usage Patterns of Research Method Groupings

For any given study, a variety of research methods are typically used to understand a dataset. We refer to groups of frequently co-occurring research methods as *method groupings*. We used a data-driven tensor decomposition approach to discover commonly used method groupings across BLS disciplines. The procedure was carried out as follows: 1) entity method by entity method co-occurrence matrices were computed by discipline, 2) co-occurrence values within each matrix were log-transformed to reduce the influence of frequently occurring method entities, 3) co-occurrence matrices for each discipline were L2 normalized to remove the effect of differing total article counts between disciplines, 4) the log-normalized co-occurrence matrices were arranged into a three-way tensor (X^N×N×D^): method entity (N = 1,218) by method entity (N=1,218) by discipline (D=15), and 5) a non-negative PARAFAC/CANDECOMP decomposition was applied to the method entity co-occurrence by discipline tensor. The CANDECOMP decomposition factorizes a given tensor into a linear combination of R rank one tensors. In the present case, the 3-way entity co-occurrence by discipline tensor can be decomposed into a sum of R rank-one tensors, referred to as *components*, as follows (the sum of outer products of three vectors):

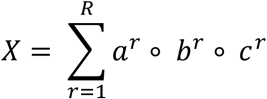

For a given component, the elements of *a* and *b* correspond to the weights of each method entity on the component (in the case of a symmetric co-occurrence matrix, *a* = *b*), and the elements of *c* correspond to the weights of each discipline on the component. As our entity co-occurrence matrices represent (log-normalized) counts, we add the additional constraint that the tensor is factorized as the additive linear sum of non-negative components. This has the benefit of enforcing sparsity on the components, and thus, increases the interpretability of the solution. There are no universally agreed upon criteria for the choice of R, the number of components. Analogous to some matrix factorization approaches (e.g., non-negative matrix factorization), the more components estimated, the finer details produced in the resulting solution. However, too many components estimated may result in redundancy, and/or modeling of noise. We chose a solution of 20 components, as this solution produced the most interpretable solution. Solutions with components around this number yielded similar solutions.

